# Single-neuron and population contributions of hippocampal LFPs to spike prediction

**DOI:** 10.64898/2026.07.14.738489

**Authors:** Ryosei Sato, Fredrich T. Sommer, Gautam Agarwal

## Abstract

Local field potentials (LFPs) contain signals generated by individual neurons and by coordinated population activity, but distinguishing these contributions remains a challenge. We examined how hippocampal LFPs at different frequencies predict single-neuron spiking during spatial navigation in male rats using two datasets. At each frequency, we assessed the spatial distribution of LFP-based prediction across the electrode array and its generalization across behavioral contexts in which a neuron remained active, but its co-active peers changed.

For pyramidal cells, spatially distributed LFP features, consistent with population-level activity, contributed primarily to spike prediction at theta (∼10 Hz) and its harmonics. In contrast, spatially localized signals, consistent with the recorded neuron’s activity, contributed predominantly at higher frequencies. Notably, gamma-band LFPs (30-80 Hz) provided comparatively little information about pyramidal-cell spiking, while distributed LFP features predicted interneuron spiking across a broader frequency range.

Together, this predictive approach separates local and distributed correlates of spiking within the LFP. In hippocampal CA1, these correlates fell into two spatiotemporal regimes: a distributed regime expressed primarily at theta frequencies and a localized regime reflecting single-neuron activity at higher frequencies.

**Significance Statement:** Brain waves reflect the activity of neuronal populations across spatiotemporal scales. We asked which features of brain waves recorded in the hippocampus predict an individual neuron’s spiking activity as a rat navigated a maze. Brain waves predicted spiking activity most accurately at two different regimes: a low-frequency, spatially distributed regime and a high-frequency, local regime. The distributed regime was concentrated in the ∼10 Hz theta band, while the local regime appeared to reflect the recorded neuron’s activity at high frequencies (>100 Hz). Surprisingly, the gamma band, often linked to neuronal communication and cell assemblies, was the weakest predictor of place cell activity. Our work provides a way to separate individual-neuron and population-level information in hippocampal LFPs.

## Introduction

Neural activity recorded in the brain is commonly categorized into two main types: spikes and local field potentials (LFPs). Beyond providing a coarse-grained measure of circuit activity, LFPs can predict the firing of individual neurons across a variety of brain regions (O’Keefe & Recce, 1993; Rasch et al., 2008; Canolty et al., 2010; Zanos et al., 2011; Ahmadi et al., 2021). Moreover, spatially distributed LFPs can predict the spiking activity of distant neurons (Canolty et al., 2010) as well as the animal’s ongoing behavior (Agarwal et al., 2014). These results suggest that LFPs reflect information encoded by spikes through the coordinated activity of neuronal populations.

In addition to the information LFPs carry about spikes, different LFP frequencies have been linked to distinct roles in neural computation. The ∼5-11 Hz theta rhythm has been associated with navigation and temporal organization of hippocampal activity (O’Keefe & Recce, 1993; Mizuseki et al., 2009), with neurons firing at progressively earlier phases of theta as the animal traverses their place fields (O’Keefe & Recce, 1993). In the gamma band (∼25–80 Hz), LFPs are thought to coordinate cell assemblies and facilitate communication within and across brain regions (Buzsáki, 2006; Lisman & Jensen, 2013; Colgin, 2015). Under these interpretations, different LFP frequency bands may carry distinct forms of information about neuronal populations. However, it remains unclear whether spike-predictive information found at different LFP frequencies reflects activity distributed across neuronal populations or localized to specific, nearby neurons.

Here, we address this question by examining how the relationship between spikes and LFPs varies across frequency and anatomical space. We adopt a framework that quantifies how well LFPs at a given frequency predict a neuron’s spiking activity. We present converging lines of evidence to determine whether, at each frequency, predictive information in the LFP is generated by the neuron being predicted, or whether it is generated by the neuron’s co-active peers.

Our results show that spike-predictive LFP signals vary by both spatiotemporal scale and cell type. We observe two regimes: a distributed regime consistent with coordinated population-level activity, and a localized regime associated with high-frequency signals near the predicted neuron. Within the distributed regime, pyramidal-cell predictions were most accurate at theta frequencies, while interneuron predictions remained accurate across a broader frequency range.

These results indicate that hippocampal LFPs contain separable population components that differentially track pyramidal-cell and interneuron activity.

## Materials and Methods

### Spike and LFP preprocessing

We analyzed data recorded from male rats running on a linear track (Mizuseki et al., 2014; Fernández-Ruiz et al., 2017). LFPs were downsampled to 1250 Hz. Spike times were binned into spike counts using a bin size of 0.8 ms (i.e., 1250 Hz).

#### Continuous Wavelet Transform (CWT)

Using MATLAB’s cwtfilterbank.m function, we pre-computed a set of complex-valued analytic Morlet wavelets 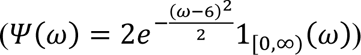 defined in terms of their angular frequency *ω*. Angular frequency (in rad/s) is converted to frequency (in Hz) by *f* = *ω*/(2*π*). We set each wavelet to 1024 samples and specified central frequencies spanning 2 to 625 Hz on a logarithmic scale using 10 voices per octave. We then convolved both the LFPs and the binned spike counts with the same wavelet bank, resulting in a complex-valued time series bandpass-filtered at each frequency.

### Spike and LFP analysis

#### Data selection

We identified “run” periods by finding indices where the rat was running on the linear track (250 cm). The rat’s position (*x*, *y*) was sampled at 39.0625 Hz. Before calculating the rat’s velocity, we low-pass filtered the position at 4 Hz using an 8th-order Butterworth filter. Then, we calculated the velocity by 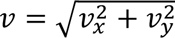 are the derivatives of and set the threshold for the “run” periods to 10% of the 99th percentile of the velocity. For the sessions analyzed, the cutoffs were 1.82 cm/s and 2.46 cm/s for the rat ec014 (n = 2 sessions) in the hc3 dataset and 5.69 cm/s in the AB3 dataset. We excluded the slower periods because they occur disproportionately at certain locations, such as the ends of the track, and are more prone to motion artifacts and sharp-wave ripples (SWRs). To isolate predictive information respectively associated with individual neurons and neuronal populations, we separated the training and testing sets according to the rat’s running direction (Fig. 1D). We trained on even trials in one running direction and tested either on odd trials in the same direction (“in-context”) or on odd trials in the other direction (“out-of-context”). We set *X* = multichannel LFPs (a matrix of size T samples x E electrodes) and *Y* = spikes (a matrix of size T samples x N neurons) to predict the latter from the former at each frequency. We excluded neurons that fired at a rate below 0.03 spikes/s during the “run” periods from the analysis, as these were poorly fit due to insufficient data. “Proximal” predictions utilized only LFPs recorded on the shank closest to the neuron of interest, while “distal” predictions utilized LFPs recorded on all other shanks. We refer to the model using all electrodes as the “full model.” The peer predictions were determined by the spiking activity of all other cells (*X* was a matrix of size T samples x (N-1) neurons).

**Fig. 1.**
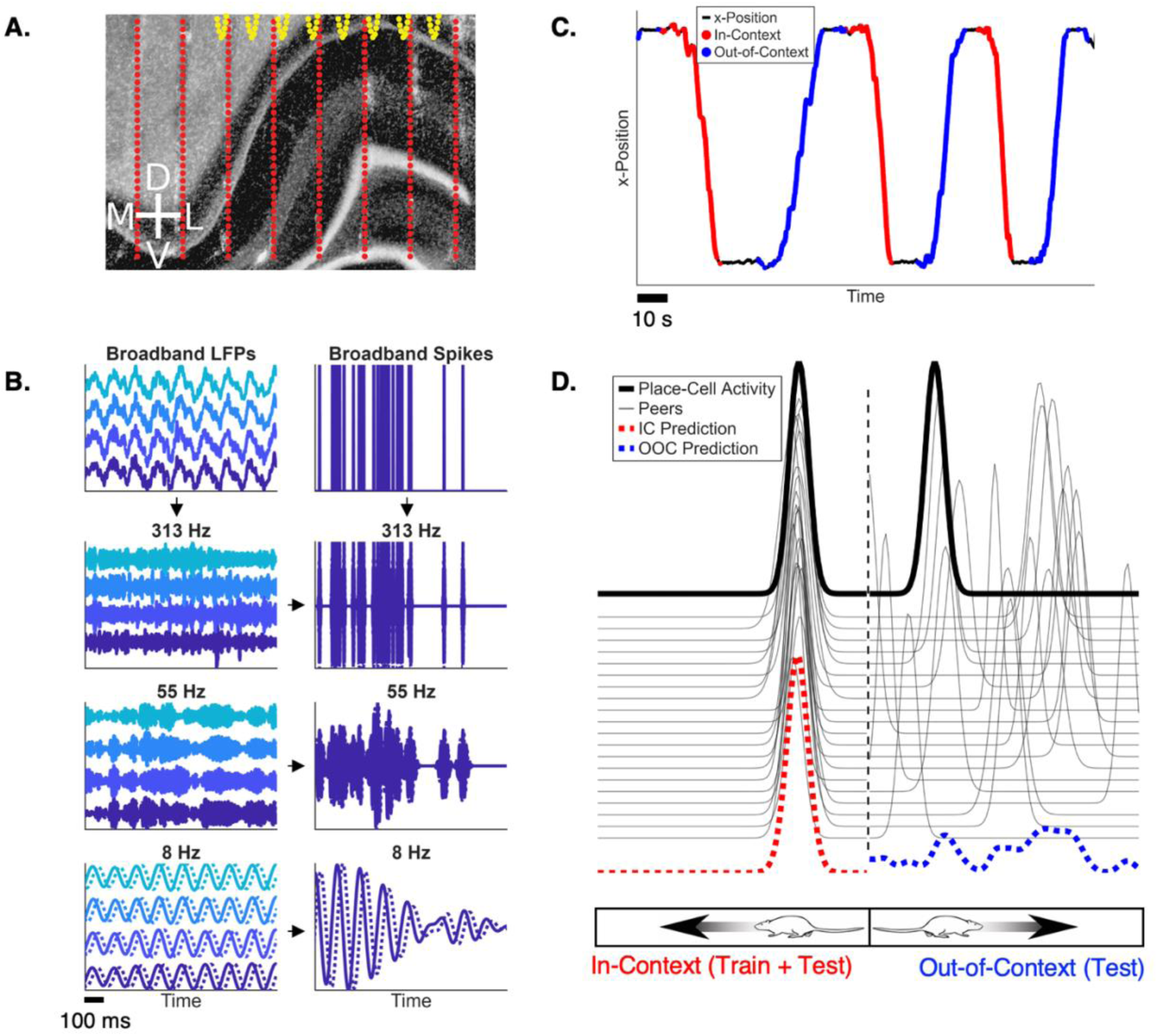
Predicting spiking activity from different LFP frequency bands. A) A 64- or 256-channel electrode array was implanted in the CA1 region of the hippocampus (panel from Agarwal et al., 2014, with permission). B) Spikes and local field potentials (LFPs), sampled at 1250 Hz, were recorded from the electrode array. Spike counts and LFPs were decomposed into frequency-specific components, which were complex-valued (imaginary components indicated using dotted lines), and separate linear models were trained at each frequency to predict spiking activity from LFPs filtered in that band. C) Rats ran on a linear track in both directions. D) Linear models were trained on data from one running direction and tested on held-out trials from the same (“in-context”, or “IC”) and opposite (“out-of-context”, or “OOC”) running directions. Differences in predictive accuracy across contexts determine whether spike information in the LFP arises from the activity of peers, which is context-specific (dotted line), or the neuron’s own activity, which is context-independent due to its stereotyped impact on the LFP.

#### Linear models

We estimated the weights w using the closed-form solution for ridge regression (Hastie, 2020). The equation is given by

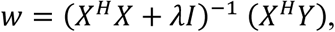

where *H* is the Hermitian operator and *I* is the identity matrix. We standardized each column of X using the z-score before solving for *w*. We solved for w at each frequency and over a grid of regularization values (*λ* = 2^0^ to 2^10^) and identified the optimal *λ* as the point at which the mean squared error (MSE) in the in-context test set was the lowest on average for all cells within the same cell type (Extended Data Fig. 2B shows the correlation coefficient rather than MSE because MSE is unbounded and obscures the influence of *λ*). To evaluate the error of the predicted (*y*^) response of a neuron relative to its actual (*y*) response (Fig. 2A), we used the complex-valued Pearson’s r,

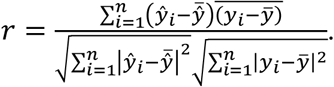

**Fig. 2.**
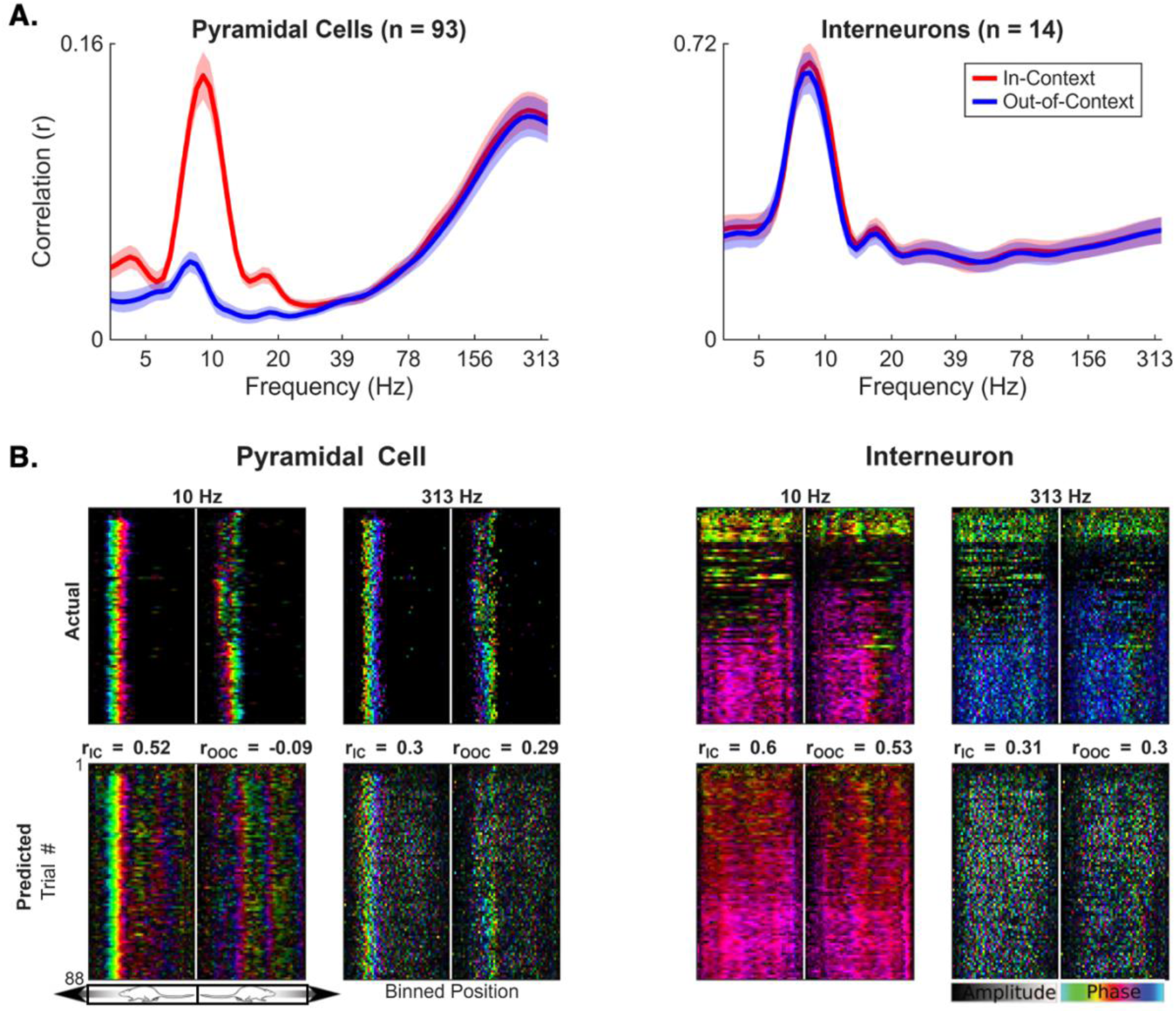
Frequency-dependent spike-LFP prediction for place cells and interneurons. **A)** Predictive accuracy (real part of the complex-valued Pearson’s correlation coefficient r, see methods) as a function of LFP frequency for pyramidal cells (left, *n* = 93 neurons from 2 sessions) and interneurons (right, *n* = 14 neurons from 2 sessions). Red (blue) lines indicate correlation coefficients calculated for in-context (out-of-context) predictions. Theta-band predictions generalize poorly across contexts for place cells but remain robust for interneurons. High-frequency predictions generalize for both cell types. Shaded regions indicate the standard error of the mean (SEM). **B)** Actual (top) and predicted (bottom) responses of a place cell and an interneuron using theta-band (10 Hz) and high-frequency (313 Hz) LFPs, as a function of position. For pyramidal cells, theta-band predictions reproduce place fields only in the training context. High-frequency predictions generalize across running directions. Colors denote the phase of spiking activity relative to the global theta rhythm. The in-context and out-of-context correlation coefficients (*r_IC_* and *r_OOC_*) for the selected neurons are noted above the predicted responses in the bottom row.

Because the predictions are expected to be phase-aligned with *Y*, we assessed model fit using the real part of *r* (ℜ(*r*)). From this point, we refer to the ℜ(*r*) as *r*.

#### Place field visualization (Fig. 2B; Extended Data Fig. 1C)

To visualize place fields at the theta band (10 Hz), we first transformed the predicted theta-band spiking activity (*Y*) (Agarwal et. al., 2014) by

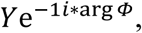

where *Φ* is the 1st principal component of the theta-band LFPs identified by principal component analysis. This transformation reveals the phase of the spikes relative to the theta wave. At higher frequencies (313 Hz), we computed

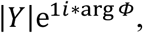

where |*Y*| is the absolute value of the high-frequency spike prediction, to visualize phase-amplitude coupling between the theta band and high-frequency LFPs. We then binned the transformed predictions at each frequency as a function of trial number and the rat’s position (Fig. 2B), or just the rat’s position (Extended Data Fig. 1C) using MATLAB’s accumarray.m with a spatial bin size of 2.5 cm on the 250 cm linear track. In each case, we visualized a place cell with place fields in both running directions.

#### Visualizing model weights

For the weights estimated by the linear models, we reshaped the one-dimensional weight vector into the two-dimensional matrix arrangement of the electrode array (Fig. 3A). To determine the distribution of weights relative to the cell body, we shifted each weight matrix so that the location of the cell is at the center of the array, padding the edges with NaNs to maintain the array dimensions. We plotted the weight magnitude, averaged across fitted neurons, as a function of relative position along the mediolateral axis (Fig. 3C; Extended Data Fig. 2D). We repeated the procedure for different frequencies and cell types.

**Fig. 3.**
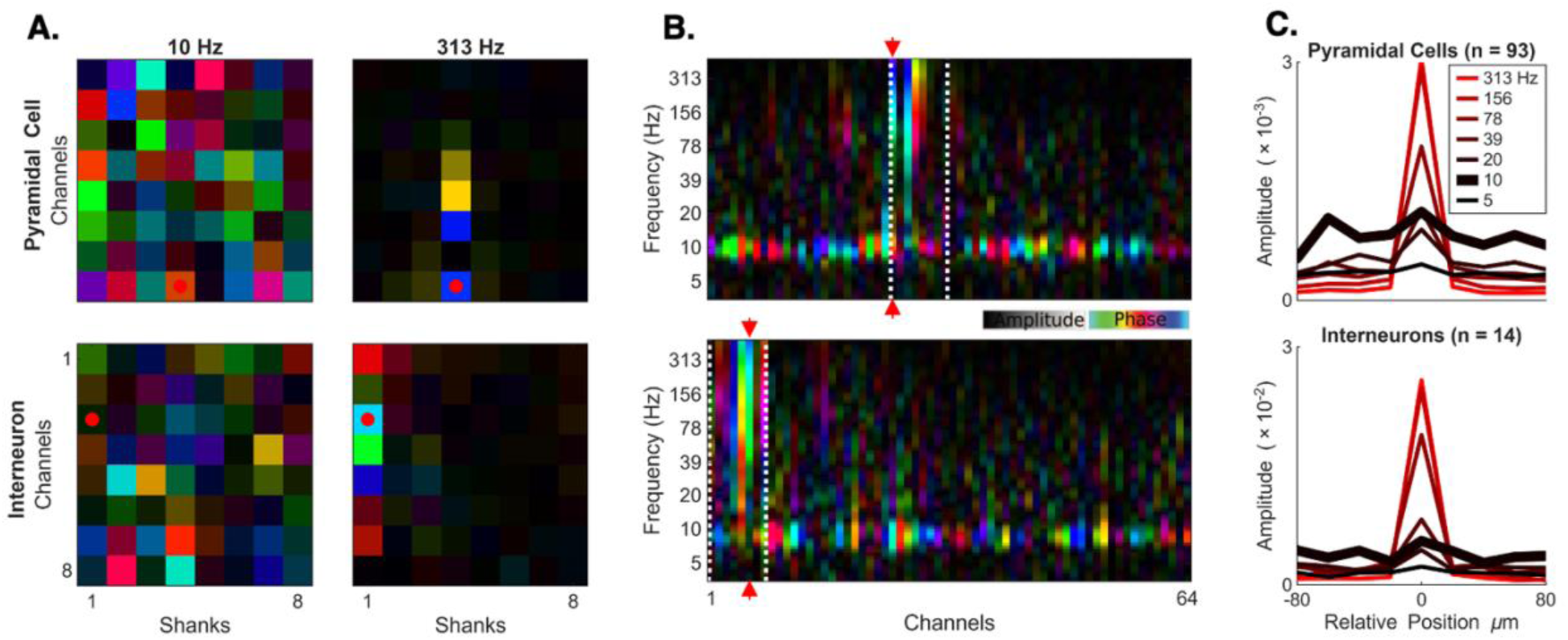
Frequency-dependent spatial structure of spike-LFP prediction. **A)** Complex-valued model weights estimated from theta-band (10 Hz) and high-frequency (313 Hz) LFPs. The weights are reshaped according to the electrode geometry in the CA1 region. Theta-band weights are broadly distributed across the array. High-frequency weights are concentrated on the shank most proximal to the cell (located at the red dot). **B)** Weights across frequencies for a pyramidal cell and an interneuron. White dotted lines demarcate the 8 electrodes belonging to the same shank as the predicted cell. At low frequencies, predictive weights are broadly distributed across electrodes; at higher frequencies, contributions become concentrated near the cell’s location (red arrow). **C)** Average magnitude of weights along the medio-lateral (M-L) axis, relative to the cell’s location. Negative position indicates more dorsal locations. Average weight amplitudes show a localized component that becomes more prominent at high frequencies (black) and a distributed component that dominates at low frequencies (red).

#### Generalized linear models (GLMs)

LFPs were first decomposed into complex-valued signals from 5-312 Hz (1 voice per octave) following the CWT preprocessing method described above. To fit GLMs with complex-valued regressors at a given frequency, we split the design matrix into real and imaginary parts. We used the softplus nonlinearity instead of the exponential function to prevent predictions of extreme values (Pillow et. al., 2008). The GLMs differed from the linear models in two respects: 1) the output (y) contained raw spike counts, rather than filtered spiking activity, and 2) while the regressors of the GLMs were filtered at a particular frequency, the GLMs’ accuracy was evaluated at all frequencies to account for any “out-of-frequency” predictions generated by the GLM nonlinearity. Therefore, we evaluated the models’ accuracy by first filtering the predicted and actual spike activity using CWT, and then calculating the correlations between them at each frequency. The GLMs were regularized with an L2 (ridge) penalty, and the optimal regularization parameter was chosen to minimize the negative log-likelihood.

#### Estimates of uncertainty

For the linear models (Fig. 2A), we performed a paired t-test comparing the accuracies (ℜ(*r*)) of models trained on theta- and gamma-band LFPs. We measured generalizability for each cell type at each frequency by regressing their models’ out-of-context accuracies on their in-context accuracies (*r_OOC_* = *β_OOC_ r_IC_*), where *β* is the best-fit slope that varies roughly between 0 (poor out-of-context generalizability) and 1 (good out-of-context generalizability) (Extended Data Fig. 1B). We estimated confidence intervals for *β* by × *SE*(*β*)), where *β* is the estimated weight, 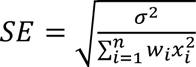 is the standard error of the weights, 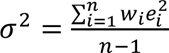 is the standard deviation of the weights, and e is the residual error, and *t_crit_* ≈ 1.992 is the critical value from the t-distribution corresponding to a two-tailed *α* = 0.05. We measured the efficacy of proximal and distal models (*r_Distal_* = *β_Distal_ r_Full_* or *r_Proximal_* = *β_Proximal_r_Full_*) for each cell by regressing their in-context accuracy against the in-context accuracy of the full model, estimating confidence intervals as described above (Fig. 4B). To assess the similarity of proximal and distal models to peer-based models (Fig. 4D), we used a linear model to predict the peer-based model accuracy at each frequency and context from the corresponding accuracies of the proximal and distal LFP models. We then performed a paired t-test to compare the best-fit weights representing the relative contributions of the proximal and distal models.

**Fig. 4.**
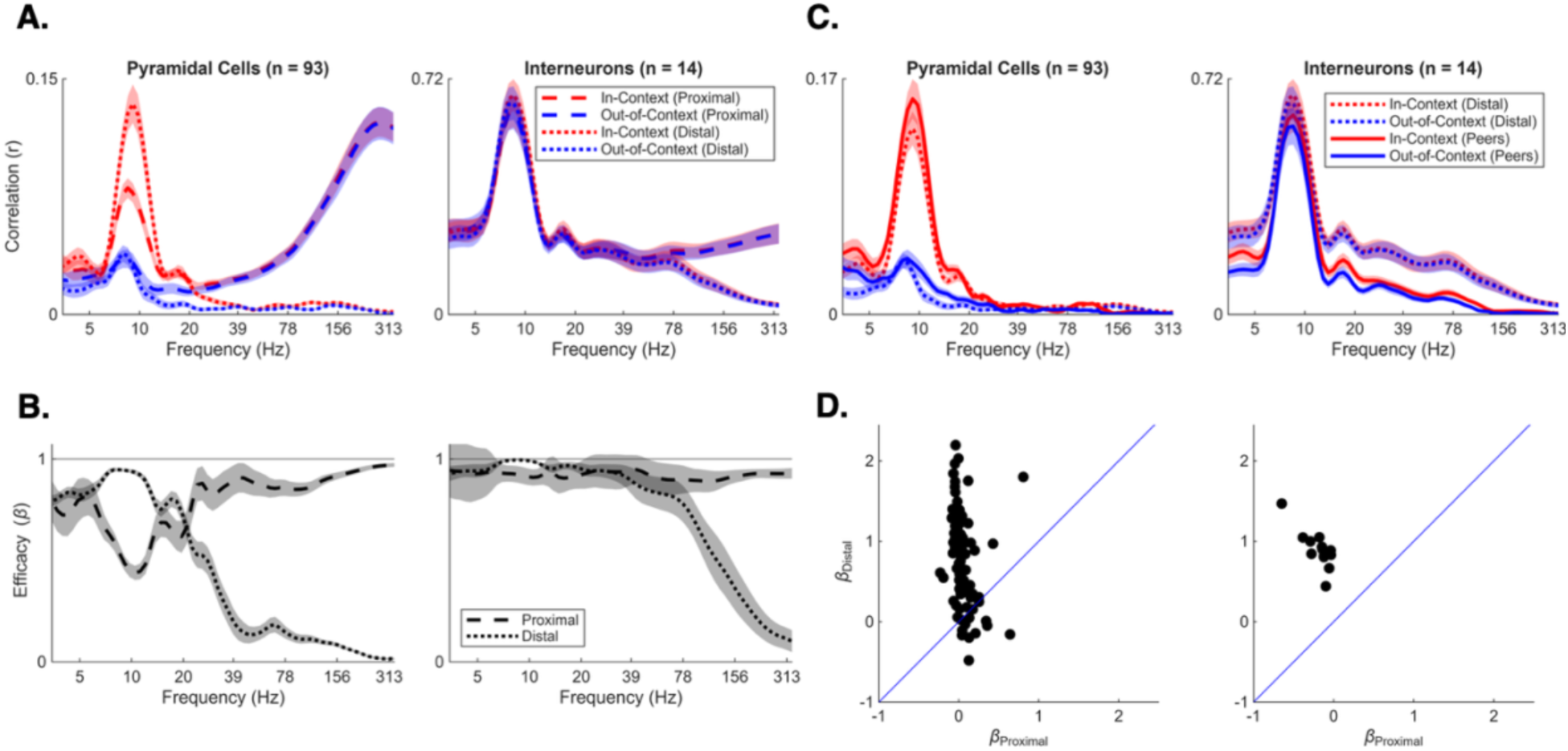
Local and distributed sources of spike prediction in LFPs. A) Predictive accuracy for models using LFPs from the shank nearest the predicted cell (“Proximal”) or from all other shanks (“Distal”). For pyramidal cells, distal theta-band (10 Hz) models outperform proximal models, with both showing decreased accuracy out-of-context. In contrast, high-frequency proximal models outperform distal models and generalize across contexts. For interneurons, proximal and distal models show similar prediction accuracy across contexts over a broader range of frequencies. B) The in-context (IC) accuracy of proximal or distal models, relative to full model accuracy (*r_Distal_* = *β_Distal_ r_Full_* or *r_Proximal_* = *β_Proximal_r_Full_*). The shaded gray indicates the 95% confidence interval (see methods). Proximal ratios increase to 1 at high frequencies for pyramidal cells; interneurons remain at around 1 for all frequencies. Distal ratios decrease at 20 Hz for pyramidal cells and 70 Hz for interneurons. C) Accuracy of models predicting pyramidal cell (left) and interneuron (right) activity using simultaneously recorded cells (peers) as regressors. Peer-based predictions closely resemble distal predictions for each cell type. Distal curves are identical to those shown in A. D) Similarity of proximal and distal models to peer-based models (see methods) for pyramidal cells (left) and interneurons (right). Each dot represents a single cell; dots above the identity line (blue) indicate greater similarity of peer-based models to distal models.

#### Power spectral analysis (Extended Data Figure 4B)

To compare the frequency responses of the different filters used in our paper and in Harris et al. (2003), we filtered pyramidal-cell spike trains using a Gaussian kernel defined by 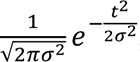, where *σ* = 25 ms, and an analytic Morlet wavelet with a central frequency of 10 Hz. Before averaging, the resulting power spectra were normalized to their maximum power within the plotted window, and the standard errors of the mean are shown.

## Results

### Frequency-resolved prediction of neural activity from multichannel LFPs

We analyzed the spiking activity of pyramidal cells and interneurons along with local field potentials (LFPs) recorded from electrode arrays with 64 or 256 channels implanted primarily in the hippocampal CA1 region (Fig. 1A). Rats ran on a 250 cm linear track in alternating directions to obtain water rewards at the ends (Fig. 1C). We focused on periods of active navigation, excluding time periods during which the rat moved slowly or was immobile, such as during reward consumption.

To examine how spike-LFP relationships vary across frequencies, we first transformed both spike counts and LFPs into complex-valued, band-limited signals using the continuous wavelet transform (CWT). We then trained separate linear models to predict spiking activity filtered at each frequency from multichannel LFPs filtered at the same frequency (Fig. 1B). Complex-valued representations naturally enable the model to learn informative phase relationships between LFPs and spiking activity (Agarwal et al., 2014; Agarwal et al., 2026).

There are two possible mechanisms through which LFPs can predict a neuron’s activity. First, LFPs may contain spatially localized signals that reflect activity specific to the predicted neuron, arising from processes near the recording site, including contributions from the action potential waveform. Second, LFPs may reflect spatially distributed activity across neuronal populations, such that a neuron’s activity can be inferred from the coordinated activity among other cells (Buzsáki, 2006; Agarwal et al., 2026). As a result, LFP-based prediction may arise from both a neuron’s own spiking and information distributed across neuronal populations.

To distinguish between these two contributions, we leveraged the direction-sensitivity of place cell responses (McNaughton et al., 1983; Fig. 1D). Linear models were trained on trials in one running direction and tested on held-out trials in both the same (“in-context”) and opposite (“out-of-context”) directions (Fig. 1D). Because population activity differs between running directions, predictions that rely on the population should generalize poorly to out-of-context directions. In contrast, predictions driven by local, neuron-specific signals should generalize well across contexts, reflecting spike-LFP relationships that are largely independent of the surrounding population state.

### Theta and high-frequency LFPs distinctly predict spike activity

Across cells, the linear models’ prediction accuracy measured by the correlation coefficient between actual and predicted responses (r, see methods) peaked at theta (∼8-10 Hz) and high (>100 Hz) frequencies, with comparatively low accuracy in the gamma range (30-80 Hz) for pyramidal cells (r*_θ_*/r*_γ_* = 0.12 ± 0.03; *H*_0_: *r_θ_* ≤ *r_γ_*, *H_A_*: *r_θ_* > *r_γ_*, *p* = 1.0 × 10^−14^) (Fig. 2A). The models predicted place fields of place cells and captured phase precession (Fig. 2B, Extended Data Fig. 1C). However, the models’ predictive accuracy depended on firing rate, cell type, and the spatial context in which a model was tested (i.e., whether models were tested in the same or opposite running direction as as the one in which they were trained) (Fig. 1D, Fig. 2A, Extended Data Fig. 1A-B).

To quantify the out-of-context (OOC) accuracy relative to the in-context (IC) accuracy, we defined a parameter *β* (0 = no accuracy; 1 = full accuracy; see methods). For pyramidal cells, the accuracy for models trained on theta-band LFPs decreased when tested out-of-context (*β_OOC_* = 0.19 ± 0.06) (Fig. 2A-B, Extended Data Fig. 1B). At higher frequencies (>100 Hz), the models maintained accurate predictions in both contexts (*β_OOC_* = 0.82 ± 0.12) (Fig. 2A-B, Extended Data Fig. 1B), suggesting that predictions depended on LFP features associated with the predicted neuron itself. In contrast, the OOC accuracy for the models predicting interneurons remained high across frequencies, including theta (*β_OOC_* = 0.95 ± 0.03) and high frequencies (*β_OOC_* = 0.98 ± 0.04 (Fig. 2A-B, Extended Data Fig. 1B). This difference in accuracy is consistent with the difference in spiking activity for the two cell types: while the spatial tuning of pyramidal cells varies by running direction (Fig. 1D), interneurons fire broadly across positions and directions and therefore are less dependent on context.

### Frequency-dependent spatial structure of spike-LFP prediction

The weights estimated by the linear models indicate which electrodes contribute predictive information about a neuron’s spiking activity in a particular frequency band. To visualize this structure, we reshaped each weight vector according to the geometry of the CA1 electrode array (Fig. 1A, yellow array). At the theta band, the models’ weights had highly variable amplitudes and phases across the full array, indicating that prediction relied on spatially distributed LFP patterns (Fig. 3A, Extended Data Fig. 2A). In contrast, at high frequencies (>100 Hz), weights with large amplitudes were confined to electrodes near the cell’s location (i.e., on the same shank), suggesting a greater contribution of local LFP features (Fig. 3A, Extended Data Fig. 2A).

To characterize this transition across frequencies, we visualized the models’ weights for all frequency bands (Fig. 3B). In line with Fig. 3A, at theta frequencies, the weights were widely distributed across electrodes. Above ∼20 Hz, weights became increasingly concentrated near the location of the predicted neuron. To test whether this relation persists across all cells, we averaged the weights after centering them on each cell’s respective location. Visualizing the mean amplitude of the weights across the medio-lateral axis revealed a broad spatial profile in the theta band and a sharp central peak at higher frequencies, with intermediate frequencies showing attributes of both (Fig. 3C, Extended Data Fig. 2C).

The distribution of the weights across frequency suggests a method for separating distributed and local sources of predictive information. We therefore trained models using either electrodes on the same shank as the predicted neuron (“Proximal”) or electrodes on all remaining shanks (“Distal”) (Fig. 3D). We refer to the model in Fig. 2A that uses all available electrodes as the “full model.” As before, we quantify the accuracy of proximal or distal models relative to the full model using a parameter *β* (0 = no accuracy; 1 = full accuracy; see methods). For pyramidal cells, distal models were more accurate in the theta band (10 Hz) than proximal models (*β_Proximal_* = 0.50 ± 0.04, *β_Distal_* = 0.95 ± 0.01) (Fig. 4A-B). At high frequencies (>100 Hz), the opposite pattern emerged: proximal models had higher accuracy than distal models (*β_Proximal_* = 0.94 ± 0.01, *β_Distal_* = 0.04 ± 0.01) (Fig. 4A-B). For interneurons, the proximal and distal models had similar accuracy across a broad range of frequencies, including theta (*β_Proximal_* = 0.92 ± 0.04, *β_Distal_* = 0.92 ± 0.03) (Fig. 4A-B). Above ∼70 Hz, distal models performed worse than proximal models (*β_Proximal_* = 0.99 ± 0.01, *β_Distal_* = 0.22 ± 0.07) (Fig. 4A-B). Together, these results indicate that low-frequency prediction of pyramidal-cell spiking relies primarily on distributed population activity, while high-frequency prediction reflects features local to the predicted neuron. Notably, this transition begins near 20 Hz for pyramidal cells and 70 Hz for interneurons, consistent with the narrower spike waveform of interneurons relative to pyramidal cells (Csicsvari et al., 1999).

### Distributed prediction is concentrated at theta frequencies

Although gamma-band activity has been proposed to organize hippocampal cell assemblies (Harris et al., 2003; Colgin et al., 2009; Buzsáki et al., 2010; Colgin, 2015), our linear models revealed little distributed predictive information about pyramidal-cell spiking in gamma-band LFPs. This result could reflect a limitation specific to the LFP—for example, that it averages out information present in the activity of single neurons. Therefore, we repeated the analysis using the activity of simultaneously recorded neurons (“peers”) rather than the LFPs as regressors. Peer-based models accurately predicted theta-band spiking activity but had low accuracies at gamma frequencies (Fig. 4C). Notably, the frequency dependence of peer-based prediction more closely resembled that of models utilizing distal LFPs than proximal LFPs, for pyramidal cells (*p* = 5.8 × 10^−21^) as well as for interneurons (*p* = 6.9 × 10^−8^) (Fig. 4D). These results support the idea that distal LFPs capture information distributed across the neuronal population.

This observation prompted us to revisit a widely cited result suggesting that the spiking activity of hippocampal pyramidal cells is best predicted from the spiking activity of peer cells at gamma timescales (Harris et al., 2003). This claim is based on the observation that a 25-ms Gaussian sliding window—nominally the period of a canonical 40 Hz gamma cycle—was optimal for smoothing peer-based responses when using them to predict a pyramidal cell’s response.

However, we found that the 25-ms Gaussian kernel acts primarily as a low-pass filter, strongly attenuating frequencies above theta (Extended Data Fig. 4). This evidence further suggests that spike-predictive information shared across hippocampal populations is concentrated at theta-related timescales rather than in the gamma band (see also Chadwick et al., 2015).

Another explanation for the poor performance of our gamma-band LFP models is that linear models lack the predictive power of approaches that better match the spike-train statistics. Specifically, generalized linear models (GLMs) (Pillow et al., 2008) define the conditional probability of a spike as a point process sampled from a Poisson distribution. Furthermore, linear models cannot predict output signals at frequencies outside of the frequency range of their regressors. As a result, they cannot detect whether gamma-band LFPs predict spiking activity at non-gamma frequencies. In contrast, GLMs rectify their outputs, thereby enabling them to generate and predict frequencies not present in their inputs. We therefore modified our approach, predicting spiking activity from bandpass-filtered LFPs using a GLM. We found that the GLM predictions resembled those of the linear model when considering spiking activity at the input signal’s frequency (Fig. 5A-B). However, high-frequency LFPs (>40 Hz) additionally predicted theta-band and broadband spiking activity in both cell types (Fig. 5A-B). For pyramidal cells, these off-frequency predictions were limited to proximal electrodes, suggesting that responses associated with the neuron, including the predicted neuron’s own spike waveform, contributed to the prediction.

**Fig. 5.**
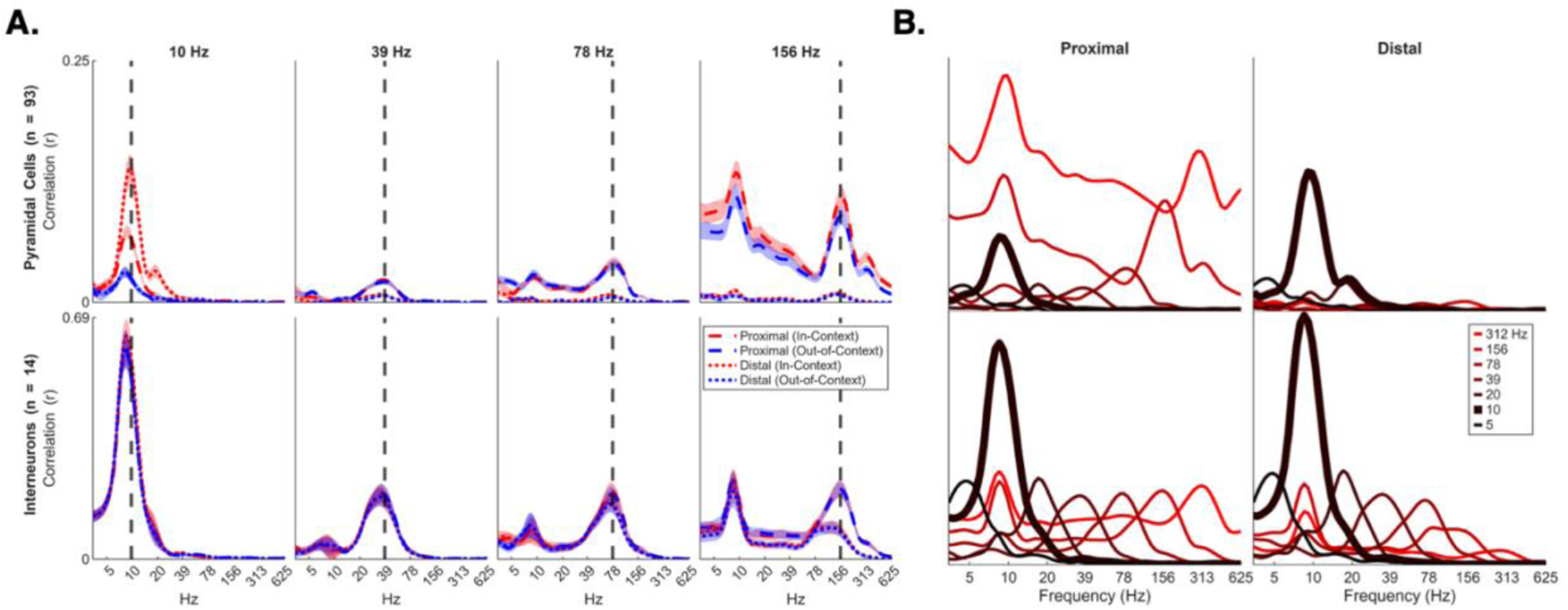
Generalized linear models (GLMs) provide cross-frequency predictions. Results are for a 64-channel array implanted in the pyramidal cell layer of CA1 (Fig. 1A, yellow array). A) Accuracy of pyramidal cell (top) and interneuron (bottom) predictions estimated by a GLM trained using LFPs filtered at different frequencies, as indicated above each column. Unlike linear models, GLMs predict spiking activity at frequencies outside the regressors’ frequency range, with prediction accuracy often peaking near theta (∼8-10 Hz). For pyramidal cells, predictions driven by regressors above theta frequencies arise primarily from LFPs recorded on the same (proximal) shank. In contrast, interneurons are similarly predicted by proximal and distal LFPs across a broader range of frequencies. B) Overlays of in-context predictions from A) for pyramidal cells and interneurons showing that regressors from a broad range of frequencies generate predictions that peak near theta. Colors from black to red (5 -> 312 Hz) indicate increasing frequency.

A third explanation is that gamma-band predictions require sampling of LFPs from deeper layers of CA1. Specifically, gamma rhythms recorded from the CA1 lac-mol layer have been shown to be phase-aligned to the spike trains of pyramidal cells (Fernández-Ruiz et al., 2021). We therefore repeated our analysis using LFPs measured from an electrode array spanning the dendritic layer of the CA1 region (Fig. 1A, red array). Nonetheless, we observed that our results were consistent with those found using the electrode array restricted to the superficial layers (Extended Data Fig. 3A-B).

## Discussion

When LFPs are related to the spiking of a neuron, care must be taken to distinguish activity associated with the recorded neuron from activity that is shared across a distributed group of neurons. In this study, we used frequency-based predictions, anatomical separation between electrodes, and generalization across behavioral contexts to separate these possible sources. These analyses revealed two regimes: a distributed, population-linked regime concentrated near theta frequencies and a localized, neuron-linked regime at higher frequencies.

In the theta band, weights estimated for pyramidal-cell prediction in our linear models were spatially distributed across the electrode array. Since CA1 place cells with similar spatial tuning are not necessarily co-localized anatomically (Slettmoen et al., 2026), we interpreted these weights as reflecting information from a distributed population of cells, potentially including cell assemblies (Hebb, 1949). Furthermore, spike prediction for pyramidal cells decreased substantially in accuracy when models were evaluated on a running direction withheld during training. In contrast to interneurons, this context dependence suggests that the linear models learned from interaction-based population activity, in which structured dynamics emerge from coupling among neurons (Freeman, 1975), rather than only phase-locking to a dominant global rhythm. By extending across frequencies and anatomical distances, our results build on earlier work that decoded position from spatially distributed LFPs (Agarwal et al., 2014) and identified shared variability among place-cell populations at theta frequencies (Agarwal et al., 2026).

At higher frequencies, we observed a different pattern. Above the theta band, the weights became increasingly concentrated near the predicted neuron, and predictions began to generalize across running directions. We interpret this transition as reflecting signals associated with the predicted neuron, including its extracellular spike waveform. This transition began at lower frequencies for pyramidal cells than for interneurons, consistent with the relatively broad extracellular waveforms of pyramidal cells. However, this transition should not be read as the frequency of the spike waveform itself, but as the frequency range at which local signals, including spike-associated and other local currents, begin to dominate over distributed predictive information.

A notable consequence of this distinction was that distributed prediction of pyramidal-cell activity was weak in the gamma band. These results do not imply that gamma is unimportant, but rather that it may have more nuanced influences: gamma may coordinate neuronal communication or support specific memory demands without strongly predicting the instantaneous firing rate of individual CA1 pyramidal cells. Within the scope of spike prediction examined here, however, the GLM, dendritic-layer analyses, and peer-based analyses provide converging support for weak gamma-band predictability.

The GLM analysis addressed whether gamma-band LFPs could predict spike-train structure outside the gamma band. Unlike the linear models, GLMs could predict theta-band and broadband components of spiking activity from gamma-band inputs, as might occur through cross-frequency coupling. Nevertheless, gamma-band LFPs remained weak predictors of pyramidal-cell spiking.

We also asked whether gamma-band prediction depended on the layers in which LFPs were sampled. Schomburg et al. (2014) and Fernández-Ruiz et al. (2021) reported strong spike-gamma phase coupling in CA1, with Fernández-Ruiz et al. emphasizing dendritic-layer LFPs. When we analyzed LFPs sampled from these layers, gamma-band activity remained a weak predictor of pyramidal-cell spiking. One explanation is that, unlike phase-locking measures, our framework asks whether LFPs predict firing-rate modulations over time, such as position-dependent modulations, rather than testing whether spikes occur at a consistent oscillatory phase.

Our peer-based analyses further motivate a reevaluation of Harris et al. (2003). That study found optimal prediction using a ∼25-ms Gaussian kernel, but the kernel is not gamma-selective and instead emphasizes theta-related and slower fluctuations. The predictive information may therefore reflect theta-related coordination rather than gamma-band oscillations per se (see also Chadwick et al., 2015).

The contrast between pyramidal cells and interneurons suggests that distributed LFP prediction does not reflect a single population signal. For pyramidal cells, distributed prediction was concentrated at theta frequencies, depended strongly on behavioral context, and was better captured by distal than proximal electrodes. This increase in distal accuracy can be explained by the greater number of distal electrodes than proximal ones, since place information becomes more accurate as the number of LFP measurement channels grows (Agarwal et al., 2014; Agarwal et al., 2026). In contrast, interneuron activity was predicted by distributed LFPs across a broader frequency range, generalized across running directions, and was similarly captured by proximal and distal electrodes. The lack of improvement with additional electrodes is consistent with the idea that theta-band interneuron activity can be predicted by the highest-variance LFP component shared across all CA1 recording sites (Agarwal et al., 2026). Thus, even within the same hippocampal network, the LFP components that predict pyramidal cells and interneurons appear to reflect distinct forms of population organization: a high-dimensional place code and a lower-dimensional interneuron-linked signal.

Our framework is related to bivariate frequency-domain measures of coupling between spikes and LFPs, such as coherence and phase-locking value. Here, we instead measure how well a multichannel LFP jointly predicts spiking activity at each frequency and provide a spatial map of the LFP features carrying information about the spikes. Unlike behavioral decoding approaches, our approach does not require specifying in advance which external variable the LFP may encode (in our case, the rat’s position). It also differs from variance-based decompositions because the most informative LFP features are not necessarily the components with the largest variance (Agarwal et al. 2026). Therefore, the method can be applied across brain regions to identify the frequencies and spatial scales at which LFPs contain information about local neuronal populations.

However, our framework is predictive rather than causal. LFPs may reflect synaptic, dendritic, and spike-related processes, and our results do not determine whether extracellular fields merely report the network state or causal influences on spike timing, such as ephaptic coupling (Anastassiou et al., 2011; Buzsáki et al., 2012). Nonetheless, spike-predictive modeling offers a principled approach for isolating LFP components that track distributed cell-assembly structure from those that primarily reflect activity near individual neurons.

## Acknowledgements

This work was supported by the Kenneth Cooke Fellowship offered by the Department of Mathematics and Statistics at Pomona College (R.S.) and startup funds from the Department of Natural Sciences at Pitzer and Scripps Colleges (G.A.). G.A. used ChatGPT (OpenAI) for language editing and manuscript organization. All content was reviewed and approved by the authors. We thank Antonio Fernández-Ruiz for helpful feedback on an earlier version of the manuscript.

## Data Availability

Data for the hc3 dataset is available at: https://crcns.org/data-sets/hc/hc-3/about-hc-3. Data for the AB3 dataset is available at: https://buzsakilab.nyumc.org/datasets/FernandezRuiz_Oliva/AB3/AB3_58_59/.

## Author Contributions

R.S. and G.A. performed research, analyzed data and wrote the paper; F.S. provided guidance and edited the paper.

## Code Availability

Code will be made available on GitHub upon publication.

---

**Extended Data Fig. 1.**
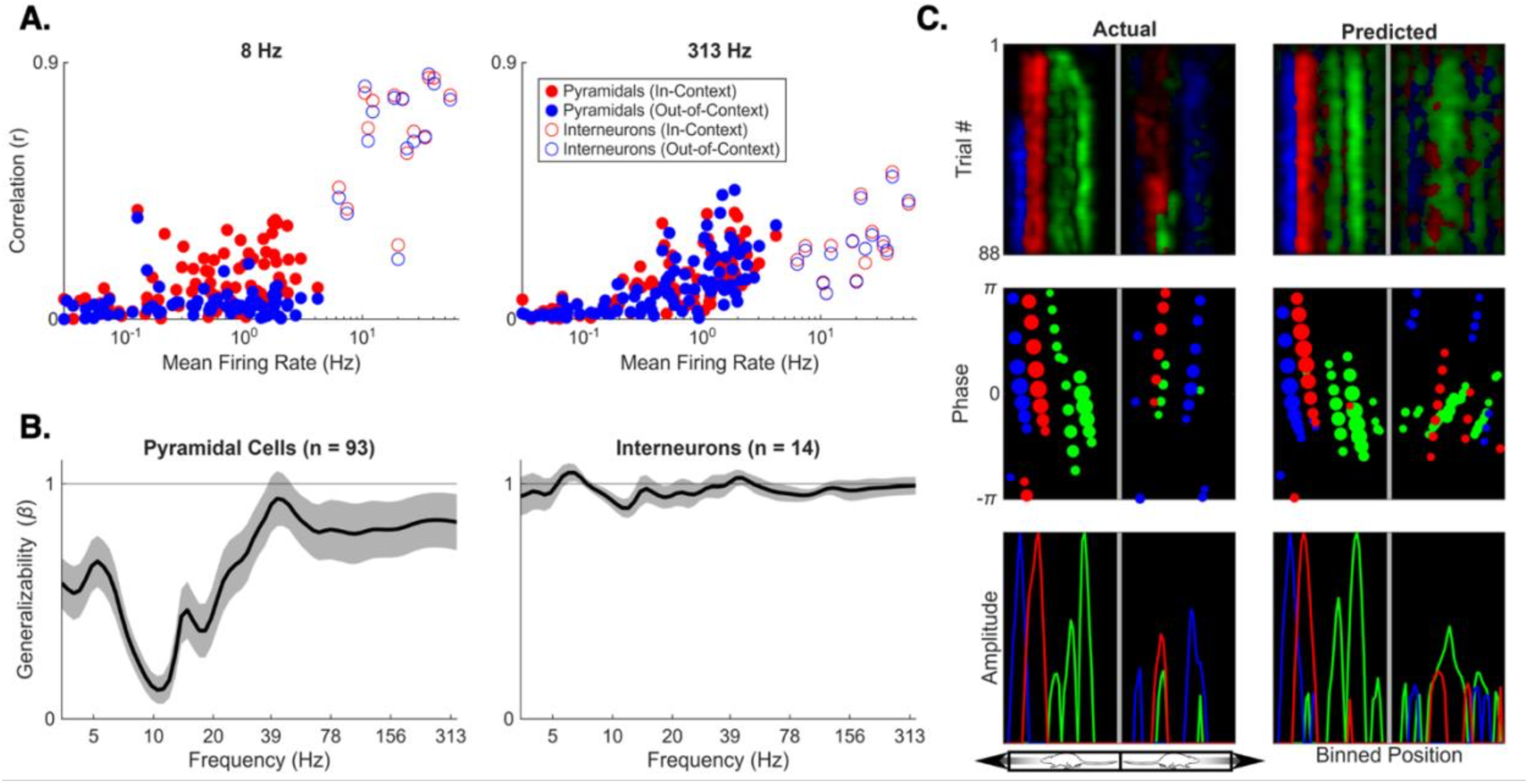
Determinants of model accuracy. **A)** Correlation coefficients as a function of the cell’s mean firing rate (spikes/second, or Hz) during running. Pyramidal cells (filled dots) and interneurons (empty dots) exhibit distinct dependencies on their firing rate, testing context, and filter frequency. **B)** Out-of-context model accuracy relative to in-context model accuracy; the shaded gray region indicates the 95% confidence interval (see methods). Pyramidal cell predictions generalize poorly across contexts in the theta band, but well at higher frequencies. Interneurons generalize well at all frequencies. **C)** Response properties of three different pyramidal cells predicted at 10 Hz. Phases are computed relative to the carrier wave (defined as PC1 of the multielectrode 10 Hz LFP) and averaged across trials. Amplitudes are calculated as the absolute value of the complex-valued firing rate, averaged across trials.

**Extended Data Fig. 2.**
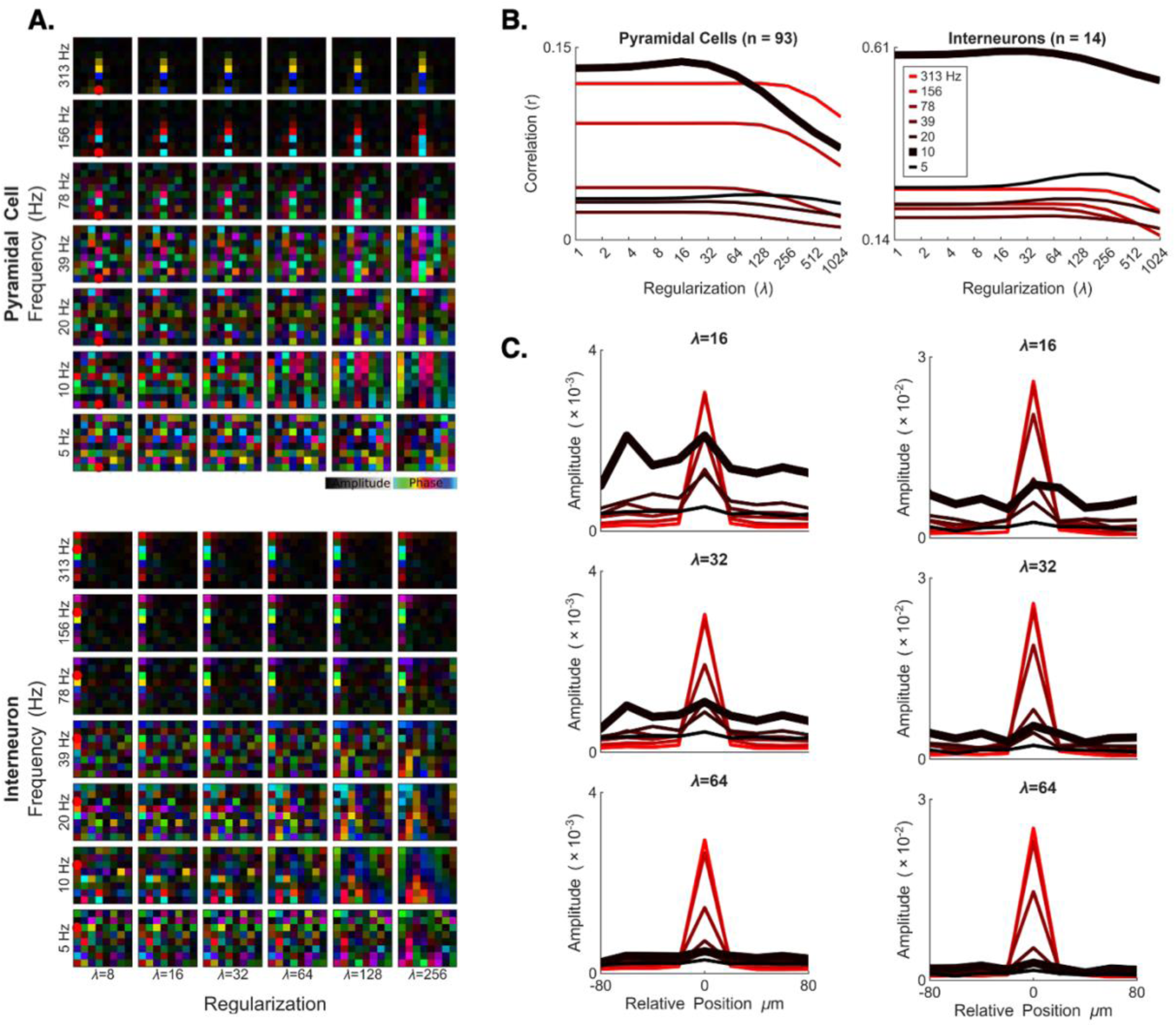
Influence of frequency and regularization on model weights and performance. **A)** Complex-valued weights for a pyramidal cell (top) and an interneuron (bottom), reshaped according to the CA1 electrode geometry. The x-axis is ordered by increasing regularization (*λ* = 8 to 256), and the y-axis is ordered by increasing frequency (5 to 313 Hz). The weight matrices show increasing smoothness as regularization increases. **B)** Prediction accuracy as a function of frequency for pyramidal cells and interneurons. Prediction accuracy peaks near *λ* = 32. **C)** Averaged weight matrices from Fig. 3C are shown for different regularization strengths (*λ* = 16, 32, and 64). Increasing regularization progressively reduces the magnitude of theta-band weights.

**Extended Data Fig. 3.**
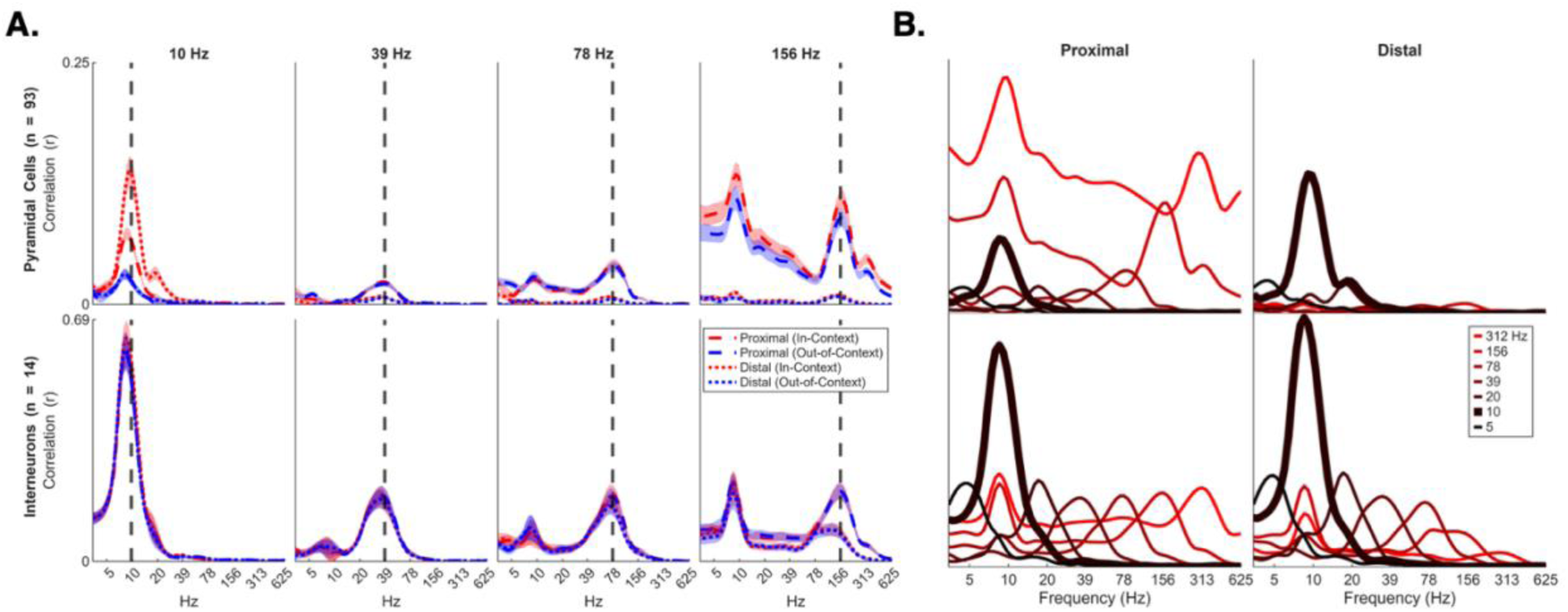
GLM predictions using LFPs recorded from a wider region of the hippocampus. Results are for a 256-channel array straddling all layers of CA1 as well as the dentate gyrus (Fig. 1A, red array). Like Fig. 4, **A)** shows the predicted accuracies (r) for pyramidal cells and interneurons across different frequencies using proximal and distal LFPs, and **B)** shows an overlay of in-context predictions across **A)**’s frequencies. The LFPs used in the prediction are sampled from the four most lateral shanks depicted in red in Fig. 1A.

**Extended Data Fig. 4.**
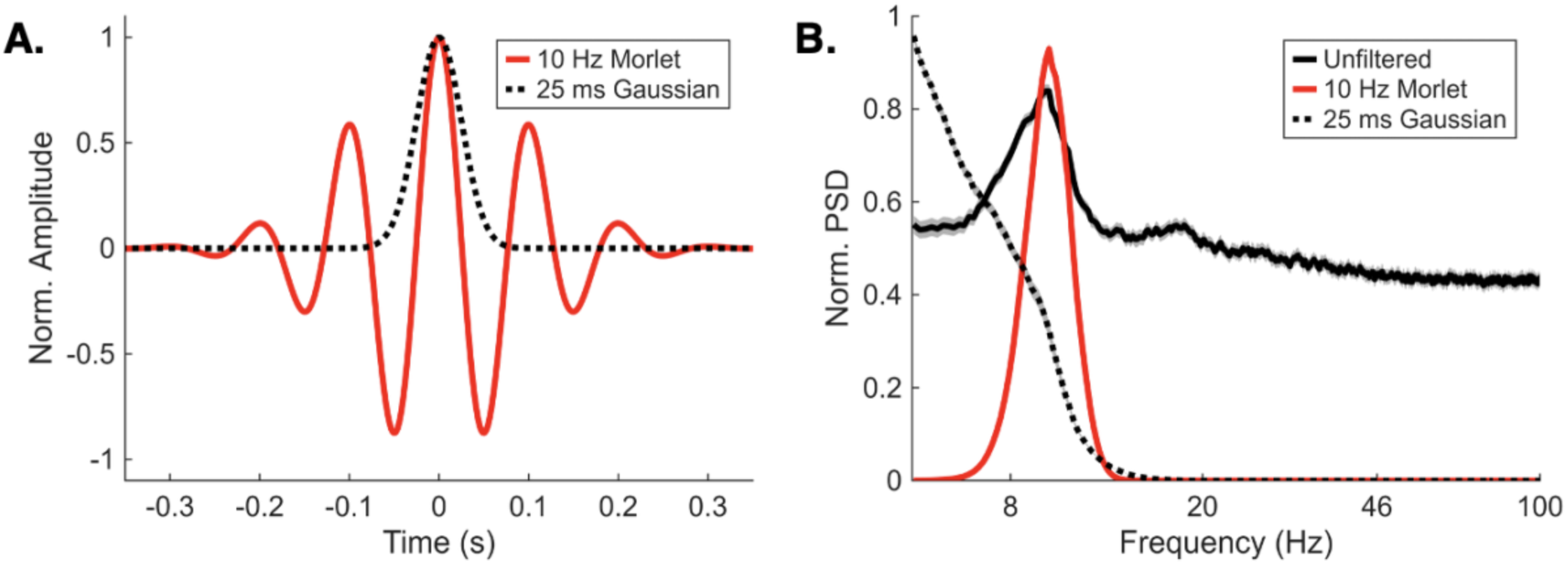
The optimal filter from Harris et al. (2003) is not a gamma-band filter. **A)** The real component of the 10 Hz analytic Morlet wavelet (red) used in the CWT of this paper, compared with the optimally predictive Gaussian kernel (black) reported by Harris et al. (2003) to correspond to gamma frequency. **B)** The power spectra of unfiltered spiking activity (black), spiking activity filtered using a 10 Hz Morlet wavelet (red), and using the optimal Gaussian filter reported by Harris et al. (2003), averaged over pyramidal cells. The power spectrum is normalized using the maximum power within the plotted range for each cell before averaging.

## References

1. Agarwal G, Akera S, Lustig B, Pastalkova E, Lee AK, Sommer FT (2026) Deciphering hippocampal place codes in weak theta rhythms. Nat Commun 17:2735.

2. Agarwal G, Stevenson IH, Berényi A, Mizuseki K, Buzsáki G, Sommer FT (2014) Spatially Distributed Local Fields in the Hippocampus Encode Rat Position. Science 344:626–630.

3. Ahmadi N, Constandinou TG, Bouganis C-S (2021) Inferring entire spiking activity from local field potentials. Sci Rep 11:19045.

4. Anastassiou CA, Perin R, Markram H, Koch C (2011) Ephaptic coupling of cortical neurons. Nat Neurosci 14:217–223.

5. Bates S, Hastie T, Tibshirani R (2024) Cross-validation: what does it estimate and how well does it do it? J Am Stat Assoc 119:1434–1445.

6. Burgess N, O’Keefe J (2011) Models of Place and Grid Cell Firing and Theta Rhythmicity. Curr Opin Neurobiol 21:734–744.

7. Buzsáki G (2006) Rhythms of the Brain. Oxford University Press. Available at: 10.1093/acprof:oso/9780195301069.001.0001 [Accessed August 29, 2025].

8. Buzsáki G (2010) Neural Syntax: Cell Assemblies, Synapsembles, and Readers. Neuron 68:362–385.

9. Buzsáki G, Anastassiou CA, Koch C (2012) The origin of extracellular fields and currents--EEG, ECoG, LFP and spikes. Nat Rev Neurosci 13:407–420.

10. Canolty RT, Ganguly K, Kennerley SW, Cadieu CF, Koepsell K, Wallis JD, Carmena JM (2010) Oscillatory phase coupling coordinates anatomically dispersed functional cell assemblies. Proceedings of the National Academy of Sciences 107:17356–17361.

11. Canolty RT, Knight RT (2010) The functional role of cross-frequency coupling. Trends Cogn Sci 14:506–515.

12. Chadwick A, van Rossum MCW, Nolan MF (2015) Independent theta phase coding accounts for CA1 population sequences and enables flexible remapping. Elife 4:e03542.

13. Colgin LL (2015) Do slow and fast gamma rhythms correspond to distinct functional states in the hippocampal network? Brain Res 1621:309–315.

14. Colgin LL, Denninger T, Fyhn M, Hafting T, Bonnevie T, Jensen O, Moser M-B, Moser EI (2009) Frequency of gamma oscillations routes flow of information in the hippocampus. Nature 462:353–357.

15. Csicsvari J, Hirase H, Czurkó A, Mamiya A, Buzsáki G (1999) Oscillatory coupling of hippocampal pyramidal cells and interneurons in the behaving Rat. J Neurosci 19:274–287.

16. Douchamps V, di Volo M, Torcini A, Battaglia D, Goutagny R (2024) Gamma oscillatory complexity conveys behavioral information in hippocampal networks. Nat Commun 15:1849.

17. Fernández-Ruiz A, Oliva A, Nagy GA, Maurer AP, Berényi A, Buzsáki G (2017) Entorhinal-CA3 Dual-Input Control of Spike Timing in the Hippocampus by Theta-Gamma Coupling. Neuron 93:1213–1226.e5.

18. Fernández-Ruiz A, Oliva A, Soula M, Rocha-Almeida F, Nagy GA, Martin-Vazquez G, Buzsáki G (2021) Gamma rhythm communication between entorhinal cortex and dentate gyrus neuronal assemblies. Science 372:eabf3119.

19. Fernandez-Ruiz A, Sirota A, Lopes-dos-Santos V, Dupret D (2023) Over and above frequency: Gamma oscillations as units of neural circuit operations. Neuron 111:936–953.

20. Freeman WJ (1975) Mass Action in the Nervous System: Examination of the Neurophysiological Basis of Adaptive Behavior Through the EEG. Academic Press.

21. Harris B, Gong P (2026) Nested spatiotemporal theta–gamma waves organize hierarchical processing across the mouse visual cortex. Nat Commun 17:2629.

22. Harris KD, Csicsvari J, Hirase H, Dragoi G, Buzsáki G (2003) Organization of cell assemblies in the hippocampus. Nature 424:552–556.

23. Hastie T (2020) Ridge Regularization: An Essential Concept in Data Science. Technometrics 62:426–433.

24. Hebb DO (1949) The organization of behavior; a neuropsychological theory. Oxford, England: Wiley.

25. Hoerl AE, Kennard RW (1970) Ridge Regression: Applications to Nonorthogonal Problems. Technometrics 12:69–82.

26. Hoerl AE, Kennard RW (n.d.) Ridge Regression: Biased Estimation for Nonorthogonal Problems.

27. Hyafil A, Giraud A-L, Fontolan L, Gutkin B (2015) Neural Cross-Frequency Coupling: Connecting Architectures, Mechanisms, and Functions. Trends Neurosci 38:725–740.

28. Kramer MA, Eden UT (2013) Assessment of cross-frequency coupling with confidence using generalized linear models. J Neurosci Methods 220:10.1016/j.jneumeth.2013.08.006.

29. Lachaux JP, Rodriguez E, Martinerie J, Varela FJ (1999) Measuring phase synchrony in brain signals. Hum Brain Mapp 8:194–208.

30. Lepage KQ, Gregoriou GG, Kramer MA, Aoi M, Gotts SJ, Eden UT, Desimone R (2013) A procedure for testing across-condition rhythmic spike-field association change. J Neurosci Methods 213:43–62.

31. Lepage KQ, Kramer MA, Eden UT (2011) The Dependence of Spike Field Coherence on Expected Intensity. Neural Comput 23:2209–2241.

32. Lisman JE, Jensen O (2013) The Theta-Gamma Neural Code. Neuron 77:1002–1016.

33. Lowet E, Sheehan DJ, Chialva U, De Oliveira Pena R, Mount RA, Xiao S, Zhou SL, Tseng H-A, Gritton H, Shroff S, Kondabolu K, Cheung C, Wang Y, Piatkevich KD, Boyden ES, Mertz J, Hasselmo ME, Rotstein HG, Han X (2023) Theta and gamma rhythmic coding through two spike output modes in the hippocampus during spatial navigation. Cell Rep 42:112906.

34. Mallory CS, Widloski J, Foster DJ (2025) The time course and organization of hippocampal replay. Science 387:541–548.

35. Markus EJ, Barnes CA, McNaughton BL, Gladden VL, Skaggs WE (1994) Spatial information content and reliability of hippocampal CA1 neurons: effects of visual input. Hippocampus 4:410–421.

36. McNaughton BL, Barnes CA, O’Keefe J (1983) The contributions of position, direction, and velocity to single unit activity in the hippocampus of freely-moving rats. Exp Brain Res 52:41–49.

37. Mizuseki K, Diba K, Pastalkova E, Teeters J, Sirota A, Buzsáki G (2014) Neurosharing: large-scale data sets (spike, LFP) recorded from the hippocampal-entorhinal system in behaving rats. Available at: https://f1000research.com/articles/3-98 [Accessed June 19, 2026].

38. Mizuseki K, Sirota A, Pastalkova E, Buzsáki G (2009) Theta Oscillations Provide Temporal Windows for Local Circuit Computation in the Entorhinal-Hippocampal Loop. Neuron 64:267–280.

39. Nadalin JK, Martinet L-E, Blackwood EB, Lo M-C, Widge AS, Cash SS, Eden UT, Kramer MA (2019) A statistical framework to assess cross-frequency coupling while accounting for confounding analysis effects Skinner FK, Colgin LL, Hyafil A, Schoffelen J-M, eds. eLife 8:e44287.

40. O’Keefe J, Recce ML (1993) Phase relationship between hippocampal place units and the EEG theta rhythm. Hippocampus 3:317–330.

41. Park M, Pillow JW (2011) Receptive Field Inference with Localized Priors. PLOS Computational Biology 7:e1002219.

42. Pillow JW, Shlens J, Paninski L, Sher A, Litke AM, Chichilnisky EJ, Simoncelli EP (2008) Spatio-temporal correlations and visual signalling in a complete neuronal population. Nature 454:995–999.

43. Rasch MJ, Gretton A, Murayama Y, Maass W, Logothetis NK (2008) Inferring Spike Trains From Local Field Potentials. Journal of Neurophysiology 99:1461–1476.

44. Rudoler JH, Herweg NA, Kahana MJ (2023) Hippocampal Theta and Episodic Memory. J Neurosci 43:613–620.

45. Schomburg EW, Fernández-Ruiz A, Mizuseki K, Berényi A, Anastassiou CA, Koch C, Buzsáki G (2014) Theta phase segregation of input-specific gamma patterns in entorhinal-hippocampal networks. Neuron 84:470–485.

46. Slettmoen T, de Jong NL, Eneqvist H, Skytøen ER, Zong W, Moser M-B, Moser EI (2026) Place cells in CA1 lack topographical organization of firing locations. Proceedings of the National Academy of Sciences 123:e2528601123.

47. Tort ABL, Kramer MA, Thorn C, Gibson DJ, Kubota Y, Graybiel AM, Kopell NJ (2008) Dynamic cross-frequency couplings of local field potential oscillations in rat striatum and hippocampus during performance of a T-maze task. Proc Natl Acad Sci U S A 105:20517–20522.

48. Wang N, Wang Y, Guo M, Wang L, Wang X, Zhu N, Yang J, Wang L, Zheng C, Ming D (2025) Dynamic gamma modulation of hippocampal place cells predominates development of theta sequences Peyrache A, Huguenard JR, eds. eLife 13:RP97334.

49. Zanos TP, Mineault PJ, Pack CC (2011) Removal of Spurious Correlations Between Spikes and Local Field Potentials. Journal of Neurophysiology 105:474–486.

50. Zhou Y, Sheremet A, Qin Y, Kennedy JP, DiCola NM, Burke SN, Maurer AP (2019) Methodological Considerations on the Use of Different Spectral Decomposition Algorithms to Study Hippocampal Rhythms. eNeuro 6:ENEURO.0142-19.2019.

51. Zutshi I, Valero M, Fernández-Ruiz A, Buzsáki G (2022) Extrinsic control and intrinsic computation in the hippocampal CA1 circuit. Neuron 110:658–673.e5.

